# Changes in pregnancy-related serum biomarkers early in gestation are associated with later development of preeclampsia

**DOI:** 10.1101/425306

**Authors:** Shiying Hao, Jin You, Lin Chen, Hui Zhao, Yujuan Huang, Le Zheng, Lu Tian, Ivana Maric, Xin Liu, Tian Li, Ylayaly K. Bianco, Virginia D. Winn, Nima Aghaeepour, Brice Gaudilliere, Martin S. Angst, Xin Zhou, Yu-Ming Li, Lihong Mo, Ronald J. Wong, Gary M. Shaw, David K. Stevenson, Harvey J. Cohen, Doff B. Mcelhinney, Karl G. Sylvester, Xuefeng B. Ling

## Abstract

**Background:** Placental protein expression plays a crucial biological role during normal and complicated pregnancies. We hypothesized that: (1) circulating pregnancy-associated, placenta-related protein levels throughout gestation reflect the uncomplicated, full-term temporal progression of human gestation, and effectively estimates gestational ages (GAs); (2) pregnancies with underlying placental pathology, such as preeclampsia (PE), are associated with disruptions in this GA estimation in early gestation; (3) malfunctions of this GA estimation can be employed to identify impending PE. In addition, to explore the underlying biology and PE etiology, we set to compare protein gestational patterns of human and mouse, using pregnant heme oxygenase-1 (HO-1) heterozygote (Het) mice, a mouse model reflecting PE-like symptoms.

**Methods:** Serum levels of circulating placenta-related proteins – leptin (LEP), chorionic somatomammotropin hormone like 1 (CSHL1), elabela (ELA), activin A, soluble fms-like tyrosine kinase 1 (sFlt-1), and placental growth factor (PlGF)– were quantified by ELISA in blood serially collected throughout human pregnancies (20 normal subjects with 66 samples, and 20 PE subjects with 61 samples). Linear multivariate analysis of the targeted serological protein levels was performed to estimate the normal GA. Logarithmic transformed mean-squared errors of GA estimations were used to identify impending PE. Then the human gestational protein patterns were compared to those in the pregnant HO-1 mice.

**Results:** An elastic net (EN)-based gestational dating model was developed (R^2^ = 0.76) and validated (R^2^ = 0.61) using the serum levels of the 6 proteins at various GAs from women with normal uncomplicated pregnancies (n = 10 for training and n = 6 for validation). In pregnancies complicated by PE (n = 14), the EN model was not (R^2^ = −0.17) associated with GA at sampling in PE. Statistically significant deviations from the normal GA EN model estimations were observed in PE-associated pregnancies between GAs of 16–30 weeks (*P* = 0.01). The EN model developed with 5 proteins (ELA excluded due to the lack of robustness of the mouse ELA essay) performed similarly on normal human (R^2^ = 0.68) and WT mouse (R^2^ = 0.85) pregnancies. Disruptions of this model were observed in both human PE-associated (human: R^2^ = 0.27) and mouse HO-1 Het (mouse: R^2^ = 0.30) pregnancies. LEP out performed sFlt-1 and PlGF in differentiating impending PE at early human and late mouse gestations.

**Conclusions:** As revealed in both human and mouse GA EN analyses, temporal serological placenta-related protein patterns are tightly regulated throughout normal human pregnancies and can be significantly disrupted in pathologic PE states. LEP changes earlier during gestation than the well-established late GA PE biomarkers (sFlt-1 and PlGF). Our HO-1 Het mouse analysis provides direct evidence of the causative action of HO-1 deficiency in LEP upregulation in a PE-like murine model. Therefore, longitudinal analyses of pregnancy-related protein patterns in sera, may not only help in the exploration of underlying PE pathophysiology but also provide better clinical utility in PE assessment.

## BACKGROUND

Placental protein expression plays a crucial biological role during normal pregnancies. The normal progression of a human pregnancy is associated with a precisely-timed transient expression of maternal and placental proteins [1, 2]. Similarly, the placenta, an endocrine gland unique to pregnancy, secretes hormones that fluctuate with respect to the gestational week of pregnancy. However, these hormones have not been useful in the development of molecular metrics to estimate GA or to phenotype complicated pregnancies prior to overt clinical manifestations of specific pathologic states like preeclampsia (PE) [3, 4], a pregnancy-related vascular disorder affecting 5–8% of all pregnancies [5, 6].

PE is thought to be a multisystem disorder of pregnancy driven by alterations in placental function and resolved by the delivery of the placenta and fetus [7]. A few pregnancy-associated, placenta-related markers have been observed having different patterns in normal pregnancies and pregnancies with PE. Chorionic somatomammotropin hormone like 1 (CSHL1; also called human placental lactogen) is selectively expressed in placental villi with an important role in regulating placental growth. Leptin (LEP) has been suggested to be involved in placental and fetal growth [8]. The relationship between LEP and PE has been discussed in a few studies [9–17]. Circulating levels of activin A, a member of the tumor growth factor protein family, can increase as early as 10–15 weeks of pregnancy in women who subsequently develop PE [18]. Elevated placental levels of angiogenic factors (soluble fms-like tyrosine kinase or sFlt-1) and decreased levels of anti-angiogenic factors (placental growth factor, PlGF) have been implicated in the pathogenesis of PE [19-25]. As such, the sFlt-1/PlGF ratio has been proposed as an index to diagnose and manage PE [26, 27]. A significant increase in sFlt-1 levels was also observed in sera of pregnant heme oxygenase (HO)-1 heterozygote (Het, HO-1^+/-^) mice, where the deficiency in HO-1 results in PE-like symptoms [28]. Recent work by Ho et al showed that PE was associated in mice with a deficiency in elabela (ELA), a placental hormone that enhances human trophoblast invasiveness in vitro [29].

In this study, we chose 6 proteins as candidates of GA estimation because they are associated with the placenta and reflect placental growth. Furthermore, levels of all 6 proteins differ in PE compared to a normal pregnancy. Previous studies found that LEP, CSHL1, and activin A are elevated early in gestation in women who subsequently develop PE [30-32]. ELA deficiency is associated with PE-like symptoms in mice [29]. The ratio of sFlt-1 and PlGF has been used clinically for diagnosing PE [27].

We hypothesized that serum levels of placenta-related proteins, LEP, CSHL1, ELA, activin A, sFlt-1, and PlGF are longitudinally regulated, and a profile of their circulating levels may collectively reflect a pregnancy-associated protein panel describing the normal progression of a term pregnancy. We further hypothesized that disruptions of this panel in early gestation are associated with placental abnormalities and an increased risk of developing PE. We sought to model the longitudinal changes in protein serum levels to estimate GA. In addition, we explored whether temporal disruptions in these profiles early in gestation are harbingers of placental pathology and subsequent PE. The model was first developed in human sera and then tested in both human and mouse sera.

## METHODS

### Study design

The study was conducted in three phases: (1) using ELISA methods to characterize the normal pattern of serum placenta-related protein levels; (2) modeling a protein-based GA estimation of normal pregnancies and identifying deviations; and (3) exploration of the protein-based GA estimation with a mouse PE model. Sera were collected in the first, second, or third trimesters during pregnancy from women who had normal uncomplicated pregnancies or a diagnosis of PE. Blood was collected at 1 to 3 time-points before week 30 of gestation and prior to a confirmatory diagnosis of PE. The GA of human was determined by ultrasound measurement. In the mouse, sera were collected from pregnant HO-1 Het or wild-type (WT) dams at 1 to 3 time-points between 7.5 to 18.5 days of gestation. The HO-1 Het mice have elevated diastolic blood pressures and plasma sFlt-1 levels during pregnancies, mimicking the PE syndrome [28]. For the human study, approval was obtained from the Stanford University Institutional Review Board. Blood was collected at Stanford University Medical Center after informed consent was obtained.

### Animal model study

For the mouse study, approval was obtained from the Institutional Animal Care and Use Committee at Stanford University. Mouse line maintenance, genotyping, and bleeding were as previously described [28].

### ELISAs

Sera from human subjects or mice were collected and measured using commercial kits specific for the human or mouse as follows: LEP (R&D System Inc., MN, USA); CSHL1 (Mybiosource, San Diego, CA, USA); ELA (Peninsula Laboratories International, Inc., San Carlos, CA, USA); activin A (R&D System Inc.); sFlt-1 (R&D System Inc.,); and PlGF (R&D System Inc.).

### Statistical analyses

Patient demographic data were analyzed using the “Epidemiological Calculator” (R epicalc package). Hypothesis testing was performed using Mann-Whitney U-tests (two-tailed). Samples collected ≥ 30 weeks of gestation or having any of the placenta-related protein measurements out of limits on the standard curves were excluded from the cohort for modeling. A 10-fold cross-validated elastic net (EN) algorithm [2, 33, 34] was used for multivariate modeling of the ELISA data. The model searches for an optimum β to minimize the least squared loss function with elastic net penalty:

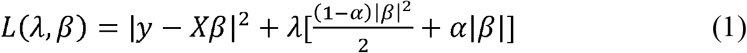

where X=(x_1_,…x_6_) is a matrix of 6 analytes, with x_j_=(x_1j_,…x_nj_)^T^, where j=1,…6. y = (y_1_,…y_n_)^T^ is the response (i.e., current GA). n is the number of samples in the training cohort. |*y – Xβ*|^2^ is the squared loss function. 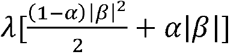 is the well-known EN penalty used for controlling the model complexity. The parameters of each penalty were controlled by α and λ. α was set to 1 and λ was set to 0.208, which maximizes the predictive value of model measured by R^2^ in the cross-validation (Additional file 1). The model is thus reduced to a lasso-regularized regression.

The mean squared error (MSE) of the GA model was used to separate PE patients from women with normal pregnancies. MSE in each woman was calculated by comparing the observed GA with the model-predicted GA. Specifically, assuming a woman had estimated GA of *ŷ_k1_*,…, *ŷ_km_*, for samples collected at the observed GA of *y_k1_*,…*y_km_*, the MSE of the model for this woman is:

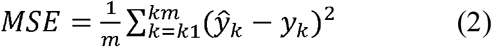

where m is the number of samples. To account for the randomness of errors, only women having 2 or more samples collected during pregnancy (i.e., m ≥ 2) were included for the calculations. Receiver operating characteristic (ROC) curves and Mann–Whitney U-tests were used to test the performance of MSE in classifying women.

The EN model in (Eq. 1) was then adjusted using 5 analytes as inputs (ELA was excluded see below). The model performance was assessed by R^2^. The role of each analyte in differentiating complicated from normal pregnancies was explored by analyzing the distribution of the concentrations at different GAs. Comparisons were made between the human and mouse to identify the common behaviors in proteins that were associated with the outcome of PE. Loess regression, Mann–Whitney U-tests, and fold changes were used for the analyses.

## RESULTS

### Samples

Forty pregnant women (20 term pregnancies, 20 with PE) receiving routine antepartum care at Stanford University Medical Center were enrolled between November 2012 and May 2016. Patient demographics are listed in Table 1. Maternal blood was collected at one, two, or three time-points during each pregnancy (at early, mid, and late-pregnancies, Fig. 1A). 10 women (4 normal pregnancies, 6 with PE) were excluded from the EN-based modeling because samples either were not collected before 30 weeks of gestation or had at least 1 protein candidate that was out of limits on its standard curve. The latter was done because outliers on the standard curve might cause distortion of continuous regression analysis. There were 30 women (16 normal pregnancies, 14 with PE) left after exclusion. Our training cohort included 10 patients who delivered at term (≥ 37 weeks GA). An independent cohort of 6 women who delivered at term and 14 women diagnosed with PE were subsequently enrolled for the validation study on normal pregnancy and the test on PE.

**Table 1.** Subject demographics.

| Characteristic | Overall Normal (n = 20) | PE (n = 20) |
| --- | --- | --- |
| **Race, No. (%)** |  |  |
| White | 20 (100) | 9 (45) |
| Asian | 0 (0) | 5 (25) |
| African-American | 0 (0) | 1 (5) |
| Other | 0 (0) | 5 (25) |
| Age, mean (SD), years | 31.9 (4.8) | 31.8 (6.0) |

| GA at delivery, mean (SD),<br>weeks | 39.5 (1.2) | 36.7 (3.3) |
| --- | --- | --- |

**Fig. 1.**
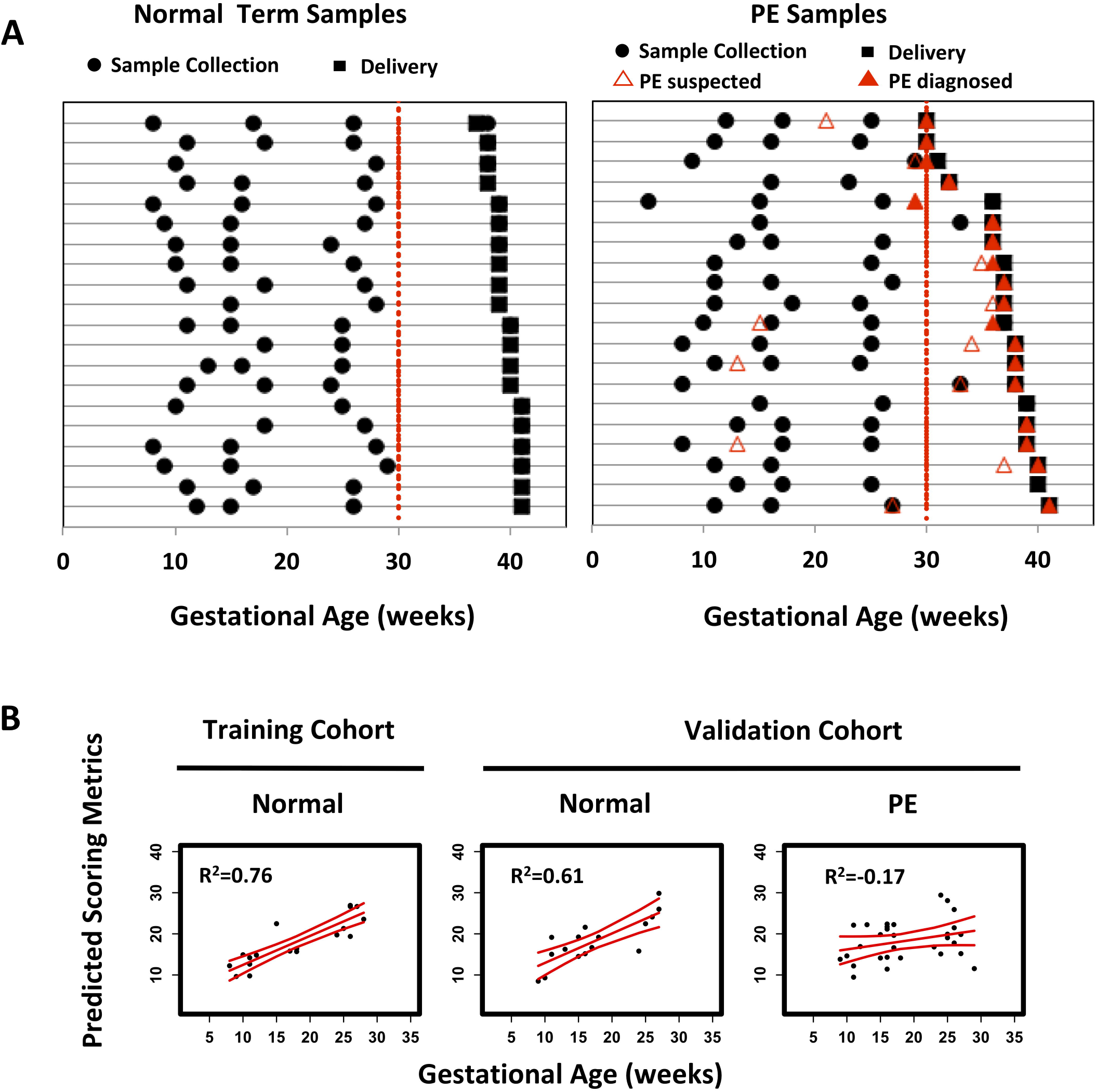
(A) Serial blood sampling from normal term and PE subjects at different GAs. Times of sample collections, infant deliveries, suspected PE, and confirmatory PE diagnoses of individual women (denoted by each row) are represented by black circles, black squares, red unfilled triangles, and red-filled triangles, respectively. (B) The EN model, developed with serial sampling analysis of 6 placenta-related proteins, dating GAs. Left panel: training cohort using sera from normal term pregnancies; middle and right panels: validation cohort using sera from normal term or PE pregnancies.

The approach was also tested with serum samples collected longitudinally from pregnant WT (n = 3 with 11 samples) and HO-1 Het (n = 4 with 15 samples) mice (Additional file 2). Each mouse had 3 or 4 samples collected at E7.5, E10.5, E14.5, and E18.5.

### A placenta-related, protein-based GA estimation of human pregnancy

We hypothesized that circulating placenta-related protein expression throughout pregnancy reflects the normal temporal progression of human gestation, and effectively serves to estimate GA. Using an EN algorithm, we developed a 6-protein model using a training cohort of 10 pregnant women that was strongly associated with GA at the time of sampling (R^2^ = 0.76, *P* = 2×10^−7^, Fig. 1B, left panel). The EN model was prospectively tested using sera serially collected from an additional 6 normal, full-term pregnant women. The EN model was found to predict GA at time of sampling in this independent normal cohort (R^2^ = 0.61, *P* = 2×10^−4^, Fig. 1B, middle panel). Univariate analyses and EN model coefficients of each protein in the model are shown in Additional file 3 and 4. Together, the analyses confirmed that there is a highly-regulated temporal pattern of protein levels in sera over the course of pregnancy (Fig. 1B).

### The placenta-related, protein-based GA estimation malfunctions in PE

Based on the above findings, we hypothesized that our EN model can identify abnormal phenotypes, such as in PE that may have an attendant disrupted placenta-related protein pattern. In contrast to the normal cohort (training R^2^ = 0.76 and testing R^2^ = 0.61, Fig. 1B), the EN model did not predict GA at time of sampling and yielded random data predictions in the PE cohort (Fig. 1B, right panel, R^2^ = −0.17, *P* = 0.2). These findings suggest that the protein-based GA estimation is disrupted in PE.

The pathogenesis of PE is complex and progresses from an asymptomatic stage, characterized by placental abnormalities during the first trimester to a symptomatic stage with proteinuria and hypertension in late gestation [35]. Our analyses revealed unique longitudinal patterns of serum protein levels of specific biomarkers (Fig. 2): LEP, CSHL1, and ELA differentiated PE from the sera of women with uncomplicated, full-term pregnancies at approximately 10 weeks of gestation, indicating that the pathogenesis of PE may arise very early in gestation. Differences in activin A begin to appear around 20 weeks and in sFlt-1 and PlGF after 25 weeks. Examination of the pattern of protein levels revealed significant protein-specific gestational windows (0–9, 10–14, 15–25, 26–33, and 27–38 weeks GA, Additional file 5 and Table 2) for each biomarker. These findings are in line with our longitudinal biomarker trending analyses (Fig. 2). Since there is a positive association [8] between maternal serum LEP concentrations and body mass index (BMI) (and consequently, gestational weight gain) during pregnancy, our analyses of LEP levels were repeated using BMI in order to normalize for serum LEP abundance. Similar findings were obtained (Additional file 6). Taken together, these data indicate that alterations in the pattern of serum protein levels of LEP, CSHL1, and ELA begin much earlier in GA than the changes in sFlt-1 (increase) and PlGF (decrease) at late GA.

**Table 2.**
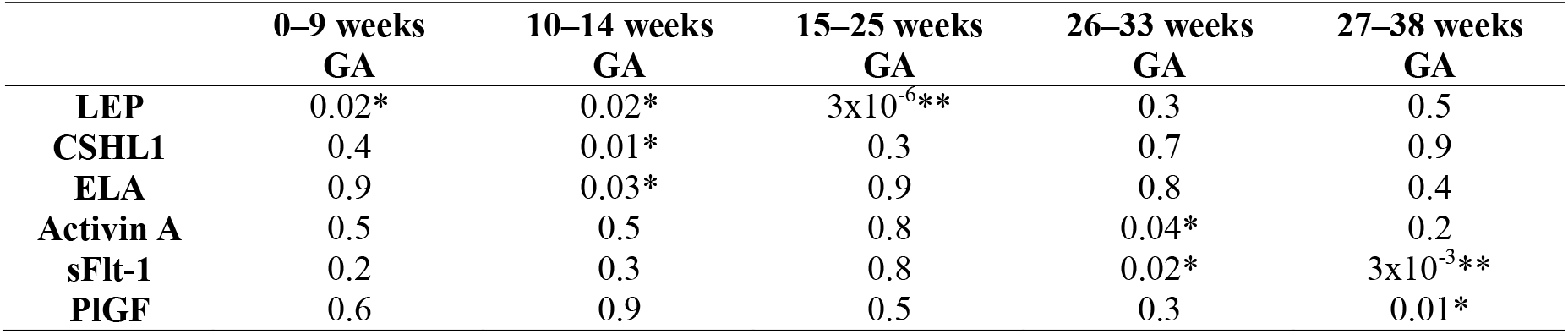
Comparisons of the serum levels of each protein between normal and PE pregnancies. Mann-Whitney U-test *P*-value was calculated. *0.005<*P*<0.05. \*\**P*<0.005.

|  | 0–9 weeks<br>GA | 10–14 weeks<br>GA | 15–25 weeks<br>GA | 26–33 weeks<br>GA | 27–38 weeks<br>GA |
| --- | --- | --- | --- | --- | --- |
| **LEP** | 0.02* | 0.02* | $3 \times 10^{-6}$** | 0.3 | 0.5 |
| **CSHL1** | 0.4 | 0.01* | 0.3 | 0.7 | 0.9 |
| **ELA** | 0.9 | 0.03* | 0.9 | 0.8 | 0.4 |
| **Activin A** | 0.5 | 0.5 | 0.8 | 0.04* | 0.2 |
| **sFlt-1** | 0.2 | 0.3 | 0.8 | 0.02* | $3 \times 10^{-3}$** |
| **PlGF** | 0.6 | 0.9 | 0.5 | 0.3 | 0.01* |

**Fig. 2.**
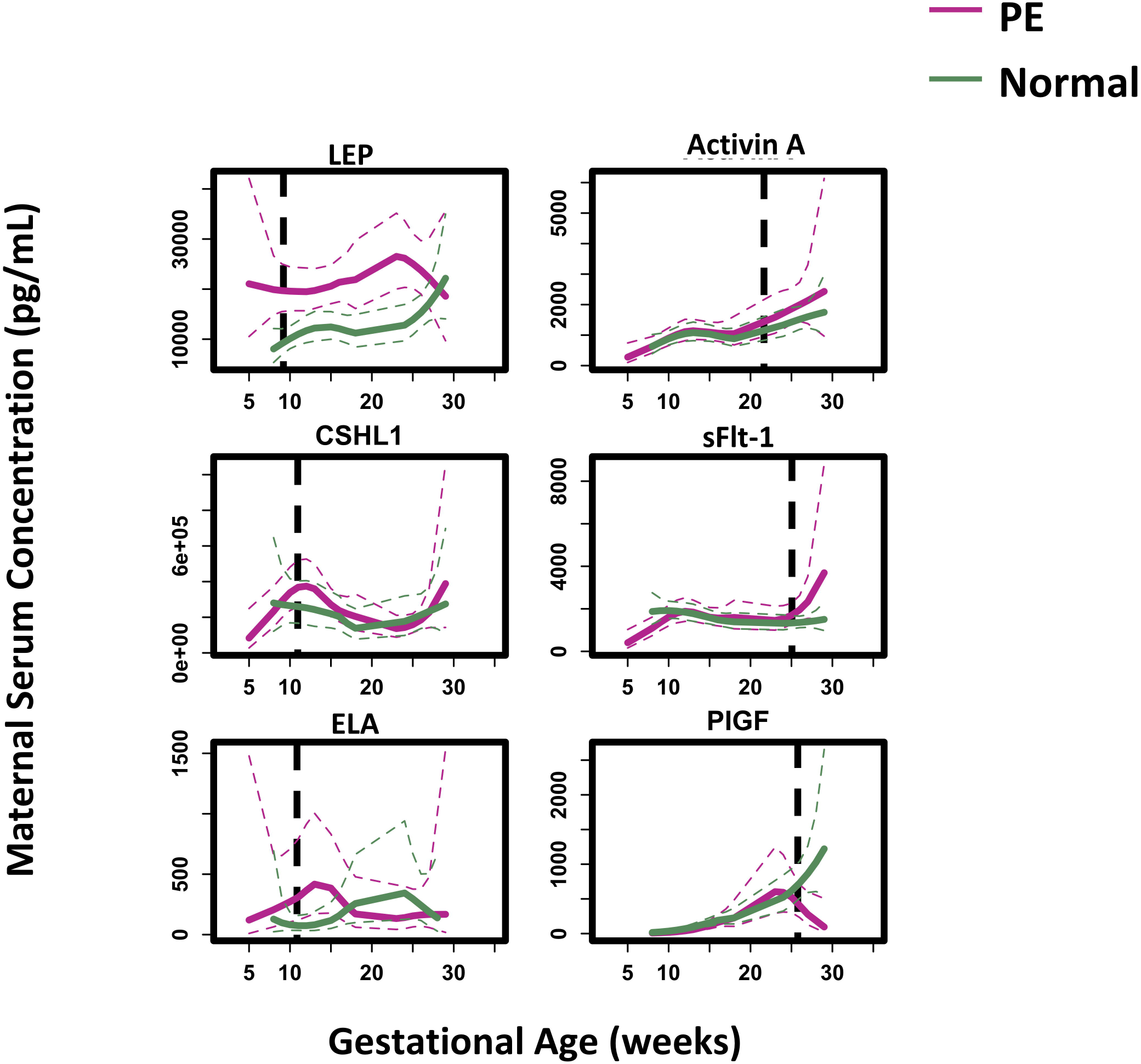
Maternal serum concentrations of 6 studied placenta-related proteins plotted as a function of the GA. Normal term pregnancies: green line. PE pregnancies: red line. Loess smooth function was applied. Color-coded dotted lines show the 95% confidence interval for each cohort.

### Disruption of the protein-based GA estimation identifies impending PE

Our placenta-related protein-based GA estimation characterizes the gestational progression of normal term pregnancies. Significant random disruptions of this normal “term” pattern were observed in women with PE. Logarithm-transformed MSE of our EN estimations were utilized to define the binary classifications to identify risk for impending PE (Fig. 3 and Additional file 7). For samples collected at 0–30 weeks GA, the MSE metric differentiated normal from PE (Mann-Whitney U-test *P* = 0.01 on the training cohort, and *P* = 0.06 on the testing cohort) with an area under the curve (AUC) of 0.88 on the training and 0.79 on the testing cohorts. An optimized cutoff value calculated on the training data yielded a positive predictive value (PPV) of 0.79 with a sensitivity of 1.00, and a negative predictive value (NPV) of 1.00 with a specificity of 0.50 on the testing data. In contrast, in a window of 16–30 weeks of gestation, performance was improved: Mann-Whitney U-test (*P* = 8×10^−3^ on the training and *P* = 0.01 on the testing) and AUC of 0.97 on the training and 1 on the testing data, PPV of 1.00 with a sensitivity of 0.88, and NPV of 0.75 with a specificity of 1.00 on the testing data. These results are better than using a single biomarker on the testing data in a window of 16–30 weeks of gestation (with AUCs of 0.53 for LEP; 0.76 for CHSL1; 0.58 for ELA; 0.53 for activin A; 0.65 for sFlt-1; and 0.65 for PlGF). Thus, our results demonstrate that significant disruptions in the protein-based GA estimation can be used to identify risk for impending PE.

**Fig. 3.**
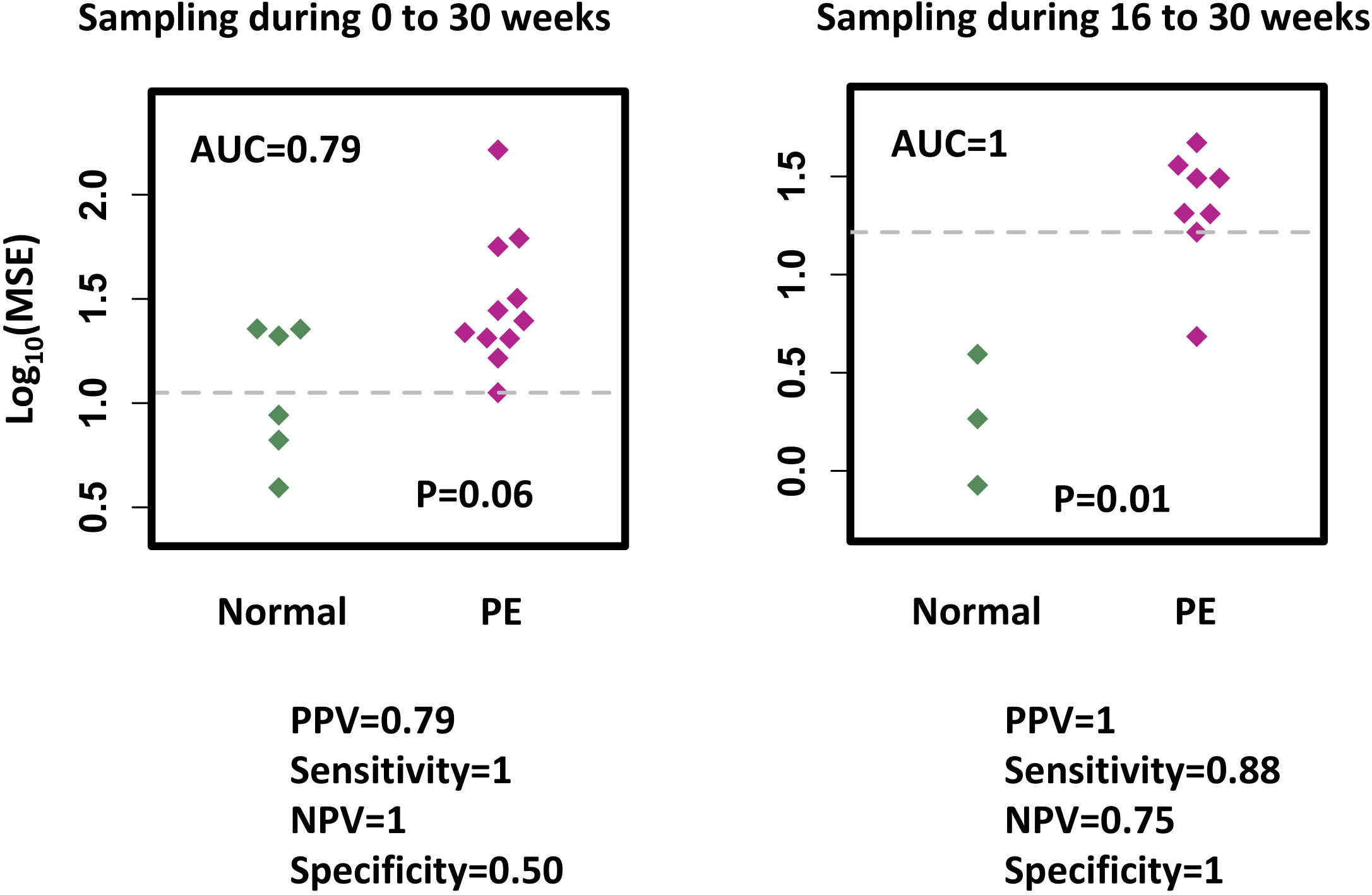
Mean squared error (MSE) of the EN model used to classify testing normal from PE. Mann-Whitney U-test *P*-value was calculated. The cut-off point (grey dotted line) shows the maximum value of the sum square of the sensitivity and 1-specificity on classification of training normal and PE cohorts at blood sampling at 0–30 weeks and 16–30 weeks GAs.

### A placenta-related, protein-based GA estimation with reduced number of features

Due to the lack of robustness of the mouse ELA ELISA essay, we tested the performance our EN-based model excluding ELA. The model had an R^2^ of 0.72 and 0.61 on the training and testing cohorts, respectively (Additional file 8). Protein pattern disruptions were observed at > 30 weeks of gestation in women with PE (R^2^ = 0.27, Additional file 8). Similar to the 6-protein model, the 5-protein model was still able to estimate GA during normal pregnancies.

### Comparative analysis of serological GA estimation between the human and a mouse model

We hypothesized that similar temporal placenta-related protein expression patterns should be conserved in mouse pregnancies, therefore, we explored our EN-based 5-protein model to normal and pregnant HO-1 Het mice, a mouse model reflecting PE-like symptoms. Model coefficients were adjusted to establish the link between GA (in days) and the targeted serum protein levels. The model had an R^2^ of 0.85 for pregnant WT and 0.30 for HO-1 Het mice with PE-like symptoms (Fig. 4 and Additional file 9). Fold changes of protein levels of HO-1 Het over normal pregnancies were calculated and then compared between the human and mouse in early (human: 5–26 weeks; mouse: 7.5–14.5 days) and late (human: 27–38 weeks; mouse: 18.5 days) gestations separately. The largest fold change was observed in LEP at late gestation of mice (Additional file 9). Unlike mice, LEP levels were elevated in women with PE in early gestation (Fig. 2, Additional file 5, and Table 2). Fold changes of sFlt-1 and PlGF in mice increased from early to late gestation. The temporal patterns of sFlt-1 in mice were similar to those in human, which decreased in early gestation and increased in late gestation of complicated pregnancies (Additional file 9). In contrast, PlGF increased in mice at late gestation, but decreased in human pregnancies after 27 weeks GA.

**Fig. 4.**
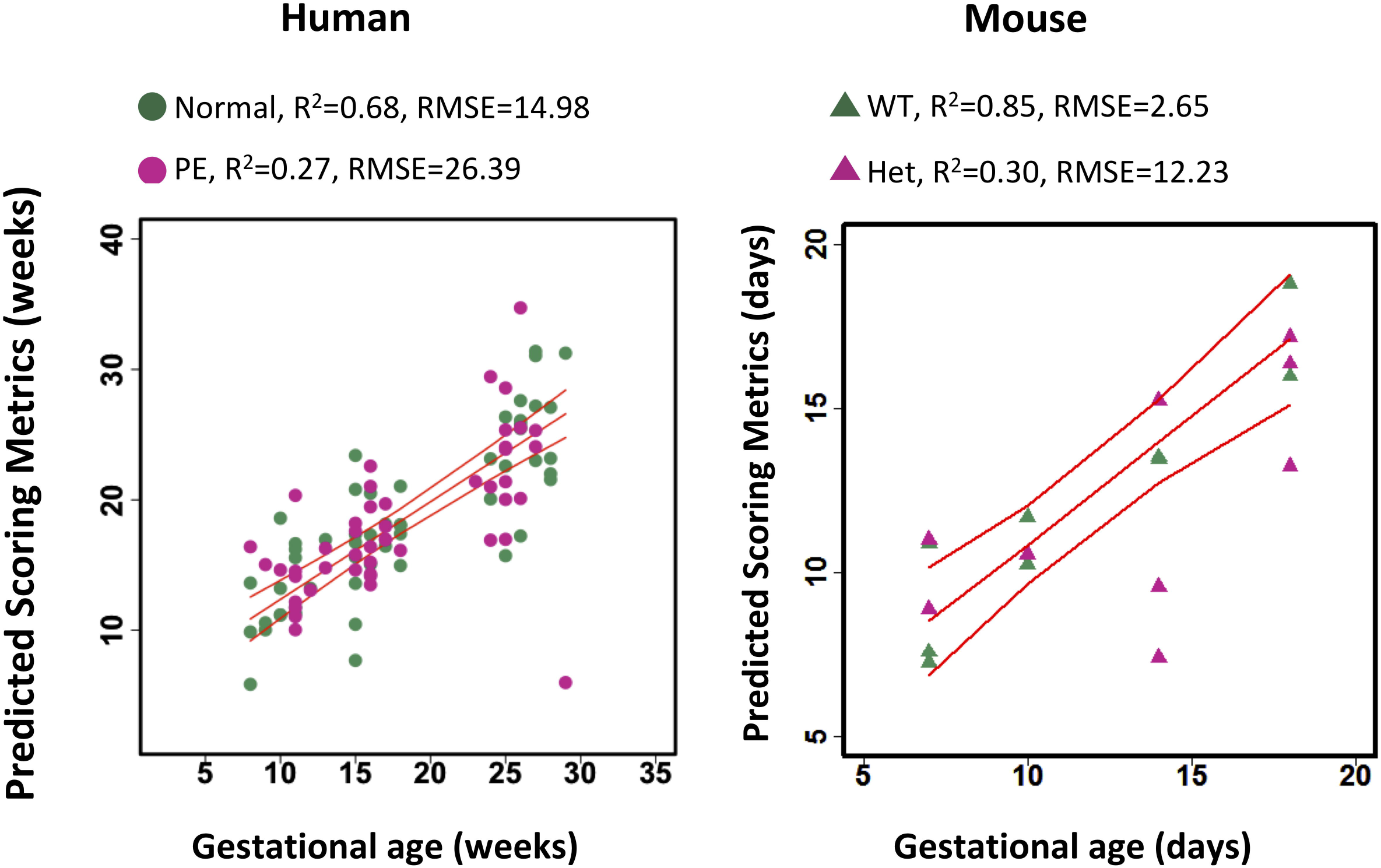
The 5-protein EN model dating GA. Left: normal human term and PE pregnancies. Right: WT and HO-1 Het mouse pregnancies (right). RMSE: root mean square error.

## DISCUSSION

The placenta plays a key role in fetal development, where cell communication occurs to support nutrition acquisition, immune adaption, and other functions of maternal-fetal interaction [36, 37]. Placental proteins are expressed in a time-dependent manner and cross-talk with other organs, such as the thyroid, pituitary, and ovary, and are necessary to ensure normal fetal development. Characterization of the temporal patterns of circulating placental proteins may serve as a basis for understanding the biology underlying both normal and pathological pregnancies. Our results support our hypothesis that multivariate modeling of the levels of circulating placental-secreted proteins, LEP, CSHL1, ELA, activin A, sFlt-1, and PlGF can be used to estimate GA during the course of a normal pregnancy, but not in women with PE. The longitudinal placental-related protein pattern in sera was also observed in pregnant WT mice but not in pregnant HO-1 Het mice.

Early diagnosis of PE remains a challenge in clinical settings. The traditional diagnosis of PE is based on the presence of maternal hypertension and proteinuria [38]. Previous transcriptomic [39-45] and proteomic [2, 46-50] profiling of normal and complicated pregnancies have identified disease-specific expression patterns and signaling networks, which suggest candidate biomarkers for possible early clinical diagnoses and for offering new biological insights. Our findings suggest that a composite placental-related protein panel from serial blood collection (for MSE calculations) may provide a diagnostic test to assess PE earlier (~10 weeks of gestation) than previously suggested by sFlt-1 and PlGF (25 weeks of gestation). Therefore, this model may offer a new investigational approach towards the understanding of placental biology during pregnancy as well as guiding innovative methods for PE diagnosis.

Our findings of serum protein levels during a normal pregnancy are consistent with those from previous studies. They are in line with the ranges reported in healthy pregnancies and have similar patterns during the pregnancy as previous results [51-59]. We found that LEP increased continuously during the first and second trimesters. Activin A remained stable between 10–20 weeks of gestation and increased late in the second trimester. sFlt-1 levels were also unchanged before 30 weeks, while PlGF progressively rose over pregnancy. We further integrated the quantitative trending information of each individual protein into a continuous regression model that expressed GA as a linear combination of the levels of proteins.

Current challenges in the management of PE include lack of early assessment and incomplete understanding of its pathogenesis and pathophysiology at early GAs. sFlt-1 and PlGF are well-established PE biomarkers [60] with clinical prognostic utilities in PE management. The ratio of sFlt-1 and PlGF has been shown to effectively differentiate PE from normal term pregnancies after 25 weeks of gestation [27]. Our findings that LEP, CSHL1, activin a, and ELA have unique serum protein signatures, starting from early to mid-gestation, are novel and that disruptions in the normal temporal placenta-related protein pattern appear at earlier GA than the conventional PE biomarkers sFlt-1 or PlGF. The presence of high levels of LEP in early gestation may signify the impending development of PE and thus serve as an early biomarker of PE. Our analyses show that LEP can differentiate PE from normal term pregnancies at <25 weeks (*P* = 3×10^−6^ at 15–25 weeks), earlier than sFlt-1, which is consistent with the previous findings of other studies [61, 62]. Given that LEP is a master regulator of energy expenditure, the observations suggest that placental insufficiency through energy imbalance is a precursor to PE that is manifested as hypertension in mid-late gestation.

Our characterization of temporal patterns of protein levels in mice provided additional support for our hypothesis. Applying our multivariate EN modeling on mouse sera we also found an association of protein levels and GA during normal mouse pregnancies, and this relationship was disrupted in pregnant HO-1 Het mice. Among the 5 proteins studied, LEP had the largest fold change in PE-like (HO-1 Het) pregnancies. The main action of leptin is in the maternal interface during the first stage of pregnancy regulating angiogenesis, growth and immunomodulation on the placenta [63-69]. Although dysregulation of leptin levels has been found correlated with the pathogenesis of various pregnancy disorders [70], including PE, the exact mechanism of action and upstream regulators remain unknown. Our characterization of the pregnant HO-1 Het mouse PE model, for the first time, provides direct evidence of the causative action of HO-1 deficiency in leptin upregulation in a PE-like murine model. This result, together with the significant differentiating power of LEP at < 25 weeks GA in human pregnancies may indicate a mechanistic role of LEP and HO-1 in the pathogenesis of PE, and deregulation of LEP as an indicator of impending PE.

We note that placenta-related proteins have distinct temporal patterns and share common characteristics between human and rodent pregnancies. sFlt-1 is upregulated in late gestation of pregnant women with PE or pregnant HO-1 Het mice. Elevated sFlt-1 levels late in gestation in mice are consistent with the findings of a previous study [28]. PlGF is upregulated after 14.5 days GA in mice while significantly down-regulated after 27 weeks GA in humans with complicated pregnancies. LEP had a significant role in identifying PE or PE-like pregnancies of both human and mice. The maximum differentiating power of LEP is achieved at late gestation (18.5 days) in mice but in early gestation (< 25 weeks) in humans. The differences in placenta-related protein patterns between humans and mice may be explained by the different placental structures (e.g. a choriovitelline placenta is initially present in mice but absent in human; trophoblast cell invasion is restricted in mice but deep in human) and different placental endocrine functions [71-73]. Despite these differences, the similar physiological features shared in human and mouse placentas, and the associations between proteins and GA in both human and mice observed in our study, show that PE-related patterns found in human are preserved in mice. It also demonstrates that studies on rodent models can be used to study the biology of human pregnancy disorders.

This study has several limitations. First, the sample sizes for our human cohorts were small, and our population lacked racial heterogeneity. Second, the time intervals of blood collections between two serial samples varied (3–31 weeks for normal, and 3–25 weeks for PE). Most samples were collected in the first or second trimester. Only 12 normal and 9 PE patients had samples collected in the third trimester. Third, serum concentrations of LEP can be influenced by maternal status [74, 75]. We addressed this through the normalization to maternal BMI (Additional file 6) and found the temporal pattern in LEP persisted. Fourth, variations in circulating protein levels could be due to the contributions from other tissues besides the placenta. Meta-analysis of PE and GA-matched uncomplicated pregnancy-associated placental gene expression patterns, including the targeted analytes of this study, has revealed similar expression trending along the gestations and differentiation between PE and normal controls [76]. Fifth, ELA was not included in the rodent analyses due to the lack of the robustness of the mouse ELISA assay. Sixth, although our protein candidates formed a panel of potential clinical utility to assess impending PE, the model robustness can be greatly improved by refinements using a sufficiently powered high time resolution cohort of sufficient powered sample size.

## CONCLUSIONS

Longitudinal EN analysis of the circulating pregnancy-associated, placenta-related protein expression throughout pregnancy revealed patterns of the normal temporal progression of human gestation which can estimate GA. The elevated MSE of the EN metric, quantifying the malfunction of the estimation, offers a potential approach to identify impending PE. The protein markers in sera shared by human and mouse and their significant associations with GA are conserved. In addition, PE-related patterns found in human are preserved in normal and HO-1 Het mice. This provides direct evidence of the causative action of HO-1 deficiency in LEP upregulation in a PE-like murine model. All of these demonstrate that the exploration of the temporal expression patterns of the placenta-related proteins in rodent models can be used to study the biology of human pregnancy disorders like PE.

With our initial placental protein-based model for PE, follow-up studies with larger, high time resolution cohorts of frequent samplings at different GA need to be performed to not only validate our current findings but also reveal additional novel serological placental proteomics patterns diagnostics of other pregnancy-related complications. Future characterization of the pregnant HO-1 Het mouse PE model may shed mechanistic insights to support HO-1 causative and leptin associated pathways as important predictors of diverse pregnancy disorders and the therapeutic target for PE intervention.

## LIST OF ABBREVIATIONS

PE: Preeclampsia
CSHL1: Chorionic somatomammotropin hormone like 1
LEP: Leptin
GA: Gestational age
sFlt-1: Soluble fms-like tyrosine kinase
PlGF: Placental growth factor
HO-1: Heme oxygenase-1
ELA: Elabela
WT: Wild-type
EN: Elastic net
SME: Mean squared error
AUC: Area under the curve
ROC: Receiver operating characteristic
PPV: Positive predictive value
NPV: Negative predictive value
BMI: Body mass index

## DECLARATIONS

### Ethics, consent and permissions

For the human study, approval was obtained from the Stanford University Institutional Review Board. Blood was collected at Stanford University Medical Center after informed consent was obtained. For the mouse study, approval was obtained from the Institutional Animal Care and Use Committee at Stanford University.

### Consent for publication

Not applicable.

### Availability of data and materials

The datasets used and/or analyzed in this study are available upon request to the corresponding author.

### Authors’ contributions

XBL, KGS, and HJC contributed to the conception and design.

JY, HZ, LC, XL, RJW, and DKS contributed to the acquisition of data.

SH, YH, HZ, LZ, LT, IM, TL, YKB, VDW, NA, BG, MSA, XZ, YML, LM, GMS, RJW, DKS, and DBM contributed to the analysis and interpretation of data.

SH and XBL drafted the manuscript.

JY, HZ, LC, YH, LZ, LT, IM, XL, TL, RJW, YKB, VDM, NA, BG, MSA, XZ, YML, LM, GMS, DKS, HJC, DBM, and KGS critically revised the manuscript.

All the authors gave final approval of the version to be submitted and agreed to be accountable for all aspects of the work.

## Acknowledgements

The authors thank colleagues at the Stanford University Pediatric Proteomics group and the March of Dimes Prematurity Research Center at Stanford University for critical discussions.

## SUPPORTING INFORMATION

**Additional file 1.** Performance of EN model with respect to α and λ in our training cohort. Left: R^2^- value of the model with respect to α when λ was set to give the minimum cross-validation mean squared error (MSE). Right: Cross-validation MSE with respect to λ when α = 1.

**Additional file 2.** Serial blood collection from pregnant WT (left) and HO-1 Het (right) mice at different GAs. Sample collection days and individual mice are represented by filled circles and lines, respectively.

**Additional file 3.** Univariate analysis of serum protein concentrations with respect to GA. Linear regression coefficients as well as 95% confidence interval and Spearman *P*-values of each protein with respect to the current GA are shown.

**Additional file 4.** Coefficients of each protein analyte in the EN model. Positive and negative values indicate positive and negative correlations, respectively, between GA and the serum protein concentrations.

**Additional file 5.** Maternal serum concentrations of the 6 placenta-related proteins plotted at different GA intervals during pregnancy. Mann-Whitney U-test *P*-values are shown.

**Additional file 6.** Maternal serum concentrations of LEP (left) and LEP normalized to body mass index (BMI) (pg/mL/kg/m^2^) (right) shown as a function of GA in normal term (red line) and PE (green line) pregnancies. Loess smooth function was applied. Color-coded dotted lines: show the 90% confidence interval for each cohort.

**Additional file 7.** The mean square error (MSE) of the GA model metric was used to classify training normal from PE subjects. Mann-Whitney U-test *P*-values are shown. The cutoff levels (grey dotted lines) show the maximum value of the sum square of sensitivity and 1-specificity on classification of normal pregnancies and pregnancies with PE.

**Additional file 8.** The EN model (α = 0.79), developed with serial sampling analysis of 5 placenta-related proteins, used for dating GAs. Left panel: shows our training cohort (normal term sera); middle and right panels show prospective testing of our EN model using normal or PE serum from serially-collected blood samples.

**Additional file 9.** Serum levels of the 5 placenta-related proteins were plotted as a function of the GA during human (left) and mouse (right) pregnancies. Loess smooth function was applied. Red: PE. Green: normal. Dotted line: 95% confidence interval.

